# Protocol development for discovery of angiogenesis inhibitors *via* automated methods using zebrafish

**DOI:** 10.1101/739854

**Authors:** Antonio Mauro, Robin Ng, Jamie Yuanjun Li, Rui Guan, Youdong Wang, Krishna Kumar Singh, Xiao-Yan Wen

**Affiliations:** Zebrafish Centre for Advanced Drug Discovery, Keenan Research Center, Li Ka Shing Knowledge Institute, St. Michael’s Hospital, Toronto, Ontario, Canada; Institute of Medical Science, University of Toronto, Toronto, Ontario, Canada; Cardiovascular Sciences Collaborative Program, University of Toronto, Toronto, Ontario, Canada; Department of Pharmacology and Toxicology, University of Toronto, Toronto, Ontario, Canada; Department of Surgery, University of Toronto, Toronto, Ontario, Canada; Department of Medical Biophysics, Schulich School of Medicine and Dentistry, University of Western Ontario, London, Ontario, Canada; Department of Medicine, University of Toronto, Toronto, Ontario, Canada; Department of Physiology, University of Toronto, Toronto, Ontario, Canada

## Abstract

Their optical clarity as larvae and embryos, small size, and high fecundity make zebrafish ideal for whole animal high throughput screening. A high-throughput drug discovery platform (HTP) has been built to perform fully automated screens of compound libraries with zebrafish embryos. A Tg(Flk1:EGFP) line, marking endothelial cell cytoplasm, was used in this work to help develop protocols and functional algorithms for the system, with the intent of screening for angiogenesis inhibitors. Indirubin 3’ Monoxime (I3M), a known angiogenesis inhibitor, was used at various concentrations to validate the protocols. Consistent with previous studies, a dose dependant inhibitory effect of I3M on angiogenesis was confirmed. The methods and protocols developed here could significantly increase the throughput of drug screens, while limiting human errors. These methods are expected to facilitate the discovery of novel anti-angiogenesis compounds and can be adapted for many other applications in which samples have a good fluorescent signal.

## 1 Introduction

The Zebrafish (*Danio rerio*) is a tropical fresh water fish belonging to the minnow family. The zebrafish has recently gained popularity as a well-managed vertebrate model for human disease. One of the main reasons for this popularity is its unique combination of optical clarity as embryos and larvae and its embryological manipulability. The optical clarity of zebrafish embryos and larvae allows for the extensive examination of the onset and progression of a pathological process *in vivo* and in real time. The breeding habits and large number of offspring per mating cycle (100-200 eggs per week) are additional positives for the zebrafish. This, together with its small size, makes it ideal for statistically significant large scale whole animal high throughput screening for chemical and genetic phenotypes. [1]

A high-throughput drug discovery platform (HTP) has been built at St. Michael’s Hospital with the goal of performing fully automated screens of compound libraries using zebrafish embryos as a model. In order to accomplish this goal, a fully functional HTP had to be established. Protocols and functional algorithms were developed for each component of the system with the intent of screening for angiogenesis inhibitors. A screen for angiogenesis inhibitors will be the pilot project for this prototypical system. Also, firmware and hardware adjustments were made to render the HTP operational and user friendly.

Transgenic Tg(Flk1:EGFP) zebrafish were used in this work. The Tg(Flk1:EGFP) line presents with a vascular endothelial specific Flk1 promoter directing EGFP expression. Flk1 is a fetal liver kinase also known as vascular endothelial growth factor receptor 2 (VEGFR-2) [2]. EGFP is an enhanced green fluorescence protein with a peak excitation wavelength at 488 nm and a maximum emission wavelength at 509 nm [3,4].

Angiogenesis is a complex multistep process that requires the tight control and coordination of endothelial cell (EC) behaviour. ECs are retained in a quiescent state in the absence of pro-angiogenic stimuli. However, low-level autocrine vascular endothelial growth factor A (VEGFA) signalling helps maintain EC homeostasis [5]. Tip cells (TCs) are selected for sprouting in the presence of high levels of exogenous pro-angiogenic factors, such as VEGFA, VEGFC, and angiopoietin 2 (ANG2), and VEGF receptor 2 (VEGFR2) or VEGFR3 signalling. Delta-like 4–Notch signalling, in contrast, laterally inhibits TC fate in adjacent ECs. [6]

Pathological angiogenesis occurs with a deregulation in homeostasis. In disorders featuring excessive angiogenesis, there is a surplus of angiogenesis activators that may be accompanied by the suppression or reduction of angiogenesis inhibitors. Cancer, one of the leading causes of death worldwide, is the most prominent disease in this category. Tumour angiogenesis has been extensively studied to characterize its mechanisms and to develop targeted and more effective therapies. Although, tumors come in many forms, angiogenesis is a process to which all tumor progression is dependent. Two distinguishing features of cancer, uncontrollable cell growth and metastasis, cannot be sustained in the absence of neovascularization [7].

Inhibition of a pro-angiogenesis pathway, such as VEGF signaling, can block angiogenesis in tumors and also change or destroy existing tumor vessels [8]. VEGF inhibitors are effective in several types of cancers, however, the benefits are transient, and the vast majority of patients who initially respond to the therapies develop resistance over time [9]. This indicates the existence of alternate pathways for cancer angiogenesis.

There have been cases in which VEGF inhibitors were found to be more effective by targeting multiple pro-angiogenesis pathways. Sunitinib, a drug approved for the treatment of renal cell carcinoma (RCC) and imatinib-resistant gastrointestina stromal tumor (GIST), is an example. This compound is a receptor tyrosine kinase (RTK) inhibitor that inhibits all VEGF and PDGF receptors, which are upregulated in clear cell RCC [10]. Thus, it is possible to tailor treatments towards specific cancers with VEGF inhibitors that also inhibit alternate pathways of angiogenesis.

This report presents the development and validation of methods that will be used to screen compound libraries with the HTP in a fully automated fashion, with the intent of initially screening for angiogenesis inhibitors. The platform configuration, experimental protocol, and novel pixel count readout is outlined.

## 2 Results

### 2.1 Platform Configuration

The fully automated high throughput drug screening platform (HTP) at St. Michael’s Hospital (SMH) was constructed by Caliper (a division of PerkinElmer) [Hopkinton, Massachusetts, USA]. The hardware and software framework was assembled by Caliper and all of the operational software was developed in house with Caliper’s iLink Pro assay development and schedule planning software. The HTP is designed to work with 96-well plates and can be modified to work with 384-well plates. The HTP contains the following components: Sciclone G3 Advanced Liquid Handler [Caliper] for compound preparation and administration, 3 x Twister II robots [Caliper] for plate transport (with transfer stations to transfer plates between robots 1 and 2 and robots 2 and 3), T-Robot thermocycler [Montreal Biotech Inc., Kirkland, Quebec, Canada], Multidrop 384 [Thermo Scientific, Hudson, New Hampshire, USA] for quick liquid dispensing of a specific liquid type over an entire plate, ELx405 plate washer [BioTek, Winooski, Vermont, USA], Synergy H1 plate reader [BioTek] to read sample fluorescence and luminescence, Image Xpress Ultra confocal microscope [Molecular Devices, Sunnyvale, California, USA], COPAS XL embryo sorter [Union Biometrica, Holliston, Massachusetts, USA] to sort and dispense embryos according to fluorescent markers, STX220 Incubator [Liconic Instruments, Somerset, New Jersey, USA], and LPX220 plate hotel [Liconic Instruments] to house compound plates. Figure 1 (Fig 1) shows a top view of a 3D CAD drawing of the HTP.

**Fig 1.**
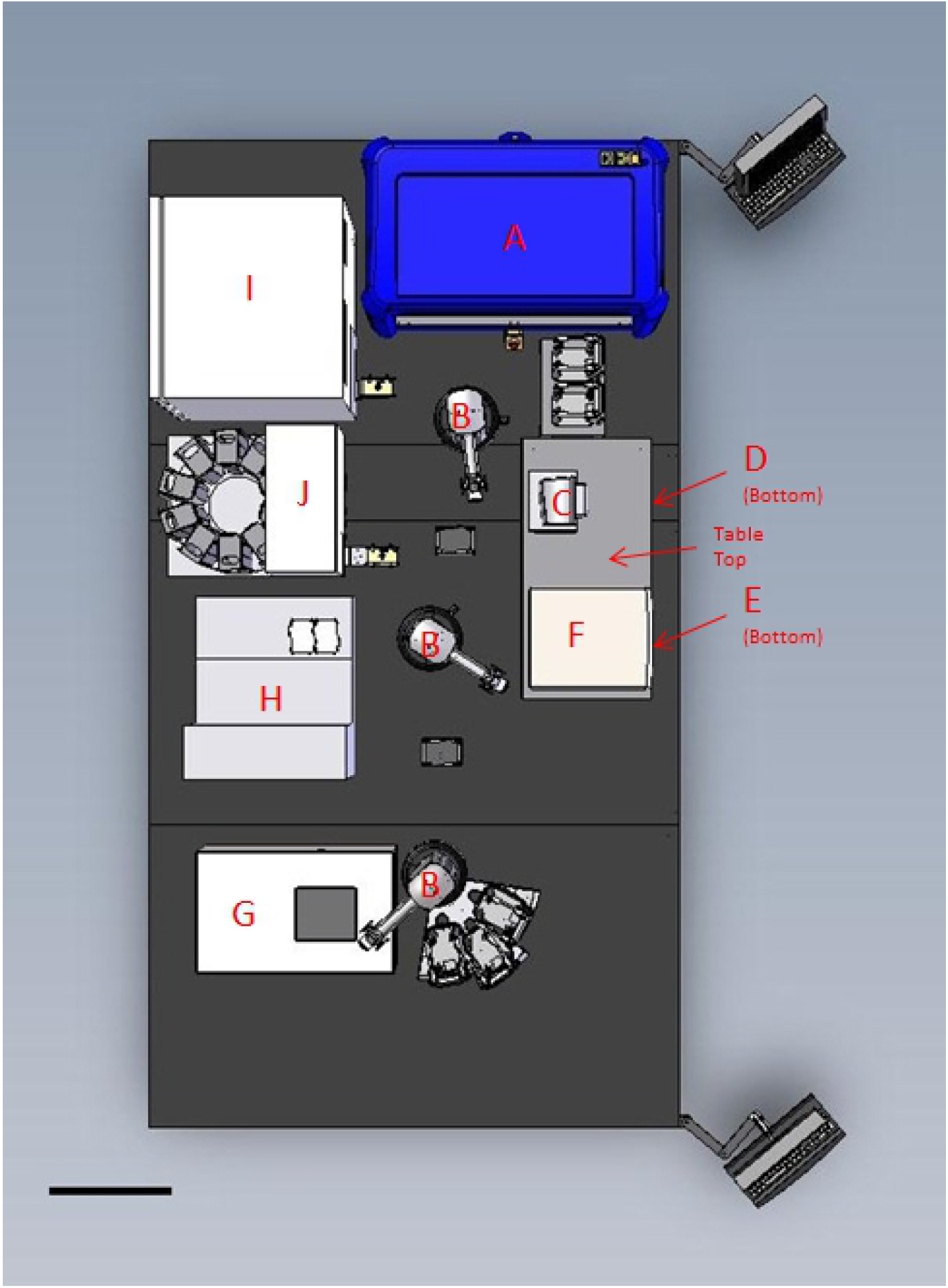
Top view of a 3D CAD drawing of the HTP. **A:** Sciclone G3 Advanced Liquid Handler. **B:** Twister II robots with transfer stations between them. **C:** T-Robot thermocycler. **D:** Multidrop 384. **E:** ELx405 plate washer. **F:** Synergy H1 plate reader. **G:** Image Xpress Ultra confocal microscope. **H:** COPAS XL embryo sorter. **I:** STX220 Incubator. **J:** LPX220 plate hotel. Scale bar is 50 cm.

### 2.2 Protocol

A drug screen for endothelial cell modulators with the HTP begins with an empty 96-well optical bottom black wall plate [Thermo Fisher Scientific, Rochester, New York, USA] in the incubator. The plate is then moved with the Twister II robots to the COPAS embryo sorter, which is filled with homozygous Tg(Flk1:EGFP) embryos at about 8 hours post fertilization (hpf) that are dispensed 1 fish/well into the 96 well plates. The optimal drop size to select 1 fish/well is 40 μL. Thus, each well contains 1 zebrafish immersed in 40 μL of E2 embryo medium as it exits the embryo dispenser. Following this, a robot brings the plate to the Multidrop 384, which adds a propylthiouracil (PTU) solution. Here, 50 μL of a PTU/E2 media solution is added per well at a 2.5x concentration of PTU (to allow for an eventual final 1.25x concentration of PTU). A PTU embryo media solution is used to block melanogenesis and leave the embryo optically clear throughout development without interfering with other processes [11]. At 10-11 hpf (beginning of the segmentation period), when primary organogenesis begins and vasculogenesis is thought to be underway [12], the plate with the samples is moved to the Sciclone along with a 96-well compound plate from the plate hotel. At this stage, 10 μL of drug from each well, at the desired concentration, is taken from the compound plate and placed into the sample plate. The compound plate is then placed back into the plate hotel and the sample plate is placed into the incubator at 32 °C. This temperature is chosen to encourage development and increase the probability that the fish will be hatched by 4 days post fertilization (dpf) [13]. The sample plate is then moved back to the Sciclone and 10 μL of 100 ppm clove oil (a zebrafish anesthetic) is placed into each well of the sample plate to immobilize the fish for imaging. From here the plate is transported to the Image Xpress confocal imager. Each well is imaged with a 4x objective. The imaging program then captures a 2D image that represents the entire length of the zebrafish. The captured images are then processed with a pixel counting program, developed as a journal in Meta Xpress [Molecular Devices, Sunnyvale, California, USA], that identifies and counts the pixels representing endothelial tissue using a lower pixel intensity threshold of 3000 on the software arbitrary scale. Figure 2 (Fig 2) is a flow chart of the HTP drug screening protocol for endothelial cell modulators. Supplementary Figure 1 (S1 Fig) is an example of the master iLink program used to instruct the HTP to execute the protocol. Figure 3 (Fig 3) is a visualization of the pixel count readout with images of a zebrafish treated with 8 μM Indirubin 3’ Monoxime (I3M), a known angiogenesis inhibitor [14], and a negative control fish treated with 0.05 % dimethyl sulfoxide (DMSO).

**Fig 2.**
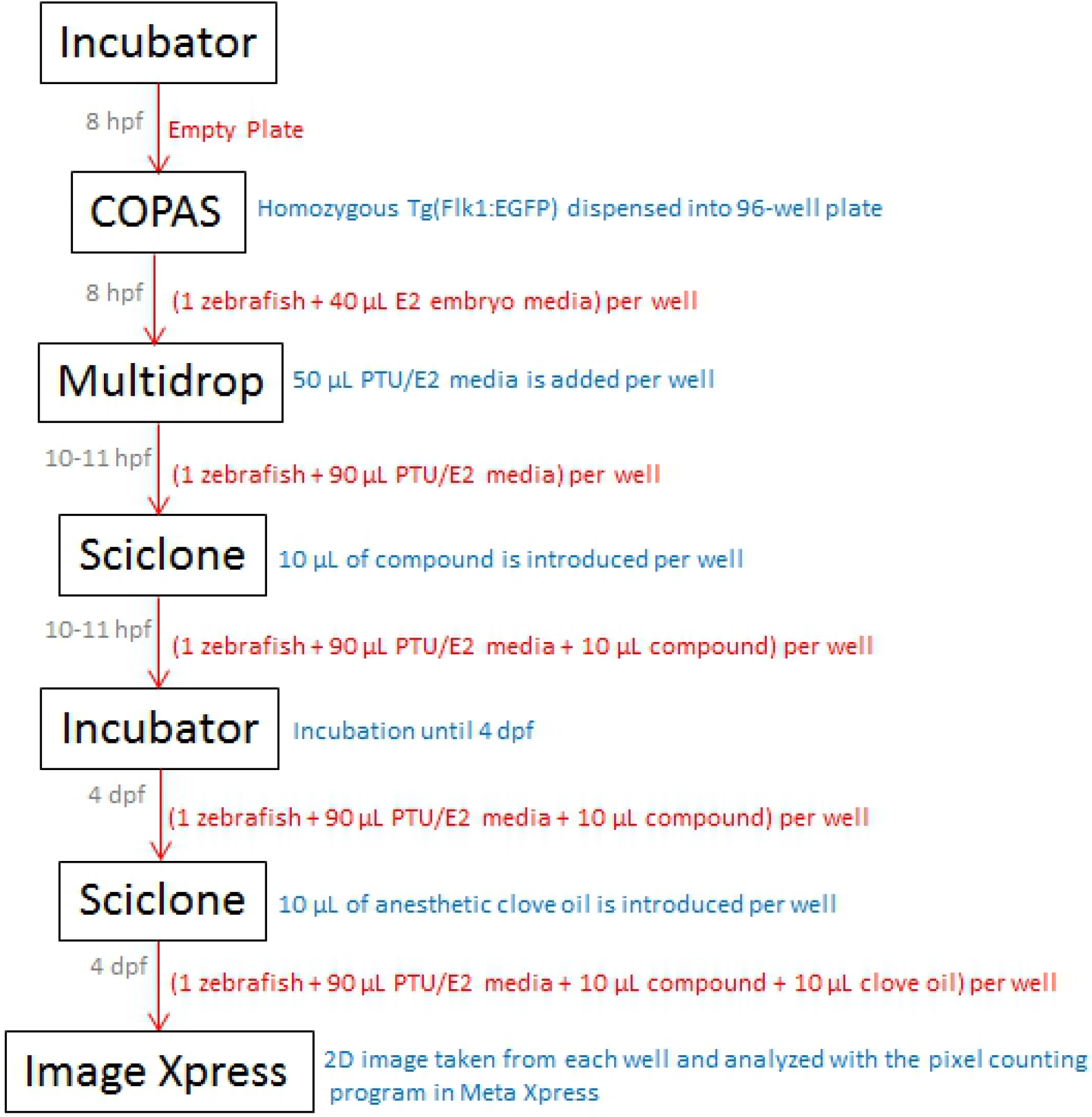
Flow chart of the HTP drug screening protocol for endothelial cell modulators. Grey font: time of plate movement. Red font: contents of plate while in transit between devices. Blue font: action taken at device.

**Fig 3.**
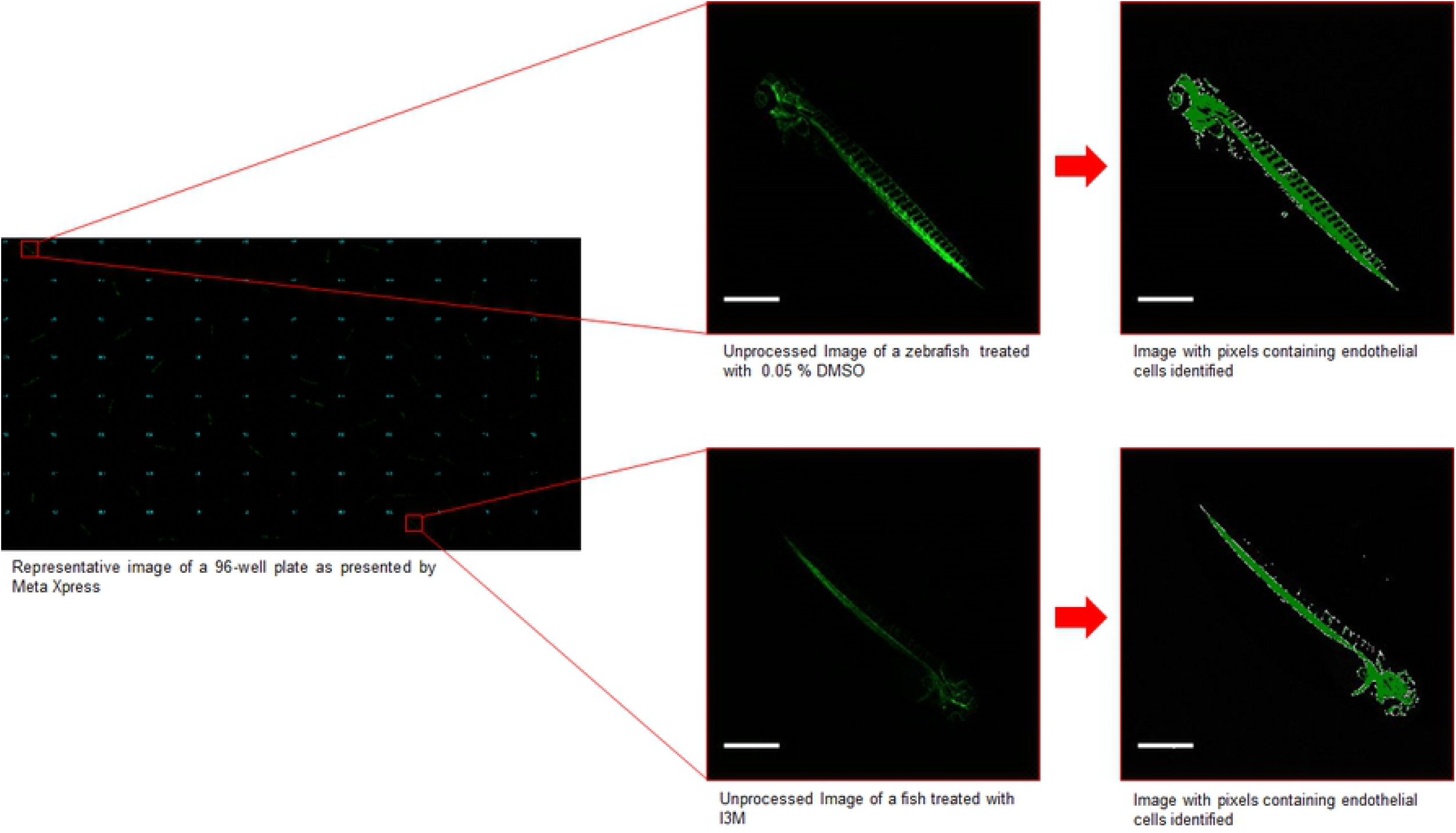
Visualization of the pixel count readout. The image on the left is a composite of 2D images from each of the wells of a 96-well plate as presented by MetaXpress. The images on the right are blown up unprocessed and processed images from a well with 0.05 % DMSO (negative control) and a well with 8 μM I3M (angiogenesis inhibitor). Scale bar is 700 μm.

### 2.3 HTP Protocol Validation

An experiment was conducted, with multiple concentrations of I3M to validate the high throughput protocol developed for the discovery of angiogenesis inhibitors. Tg(flk1:EGFP) zebrafish were distributed 1 fish/well in a 96-well plate, with two columns designated for each concentration. The concentrations of I3M were 0 μM (0.05 % DMSO control), 1 μM, 2 μM, 4 μM, 8 μM, and 16 μM (Fig 4; N = 16/group in triplicate). Values under 4000 pixels were assumed to be dead embryos and were excluded. Distorted values due to poor images attributed to moving fish were also removed. After an ANOVA with a Tukey’s post-hoc analysis, it was found that the concentrations of 8 μM and 16 μM yielded significantly (P < 0.05) lower pixel counts when compared to the control group, as shown in Figure 4 (Fig 4). I3M was shown to function in a dose dependent manner. These results were in accordance with previous literature [14], thus validating the protocol. Also, having values for mortality in each concentration group, it was possible to construct a plot representing survival (Fig 5).

**Fig 4.**
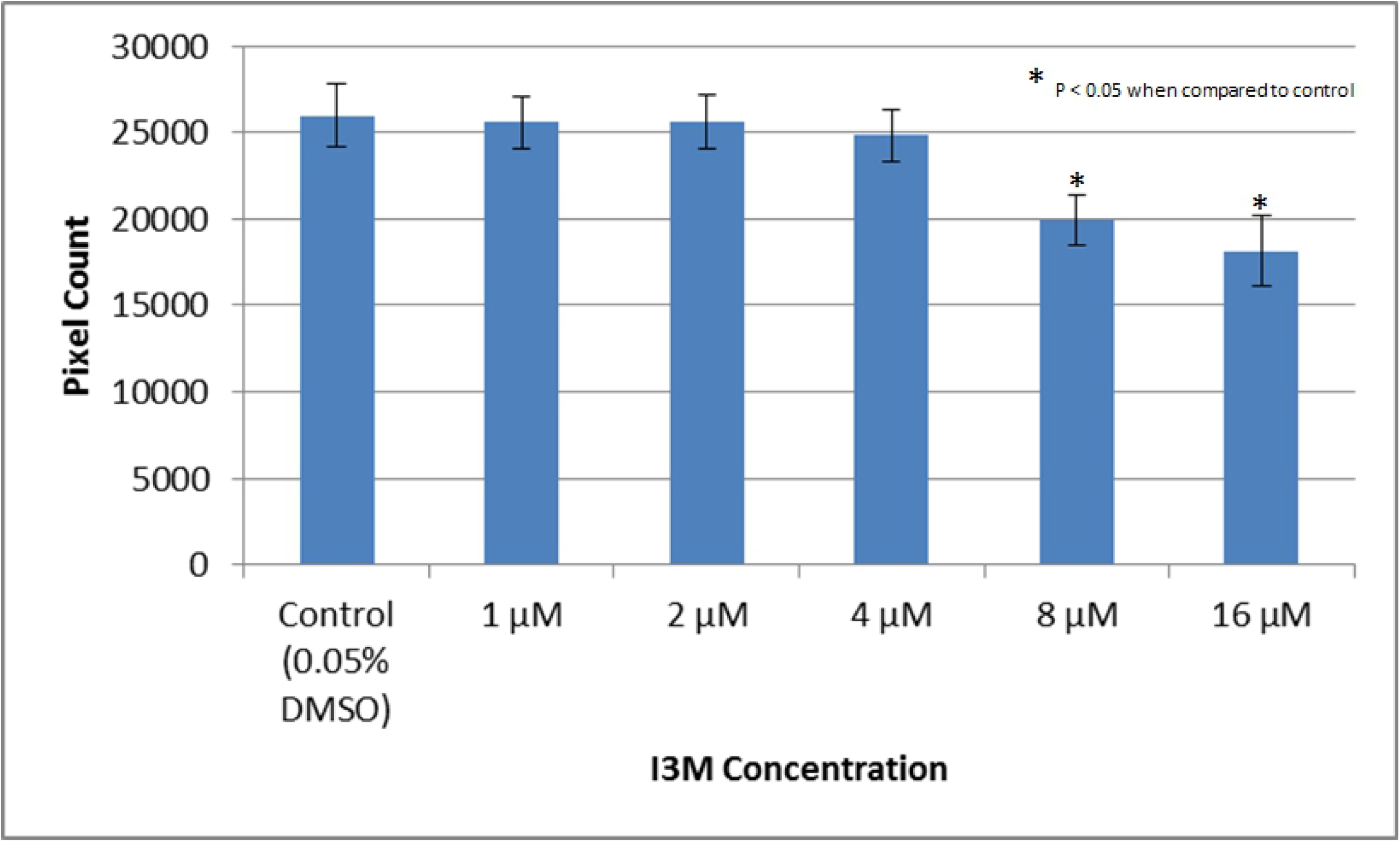
Pixel count readout results that validate the high throughput protocol developed for the discovery of angiogenesis inhibitors. Tg(flk1:EGFP) zebrafish were treated with 0.05% DMSO and compared to Tg(flk1:EGFP) zebrafish treated with various concentrations of I3M to validate protocol. The experiment was repeated in triplicate. The error bars represent the standard deviation of the means from the three experiments.

**Fig 5.**
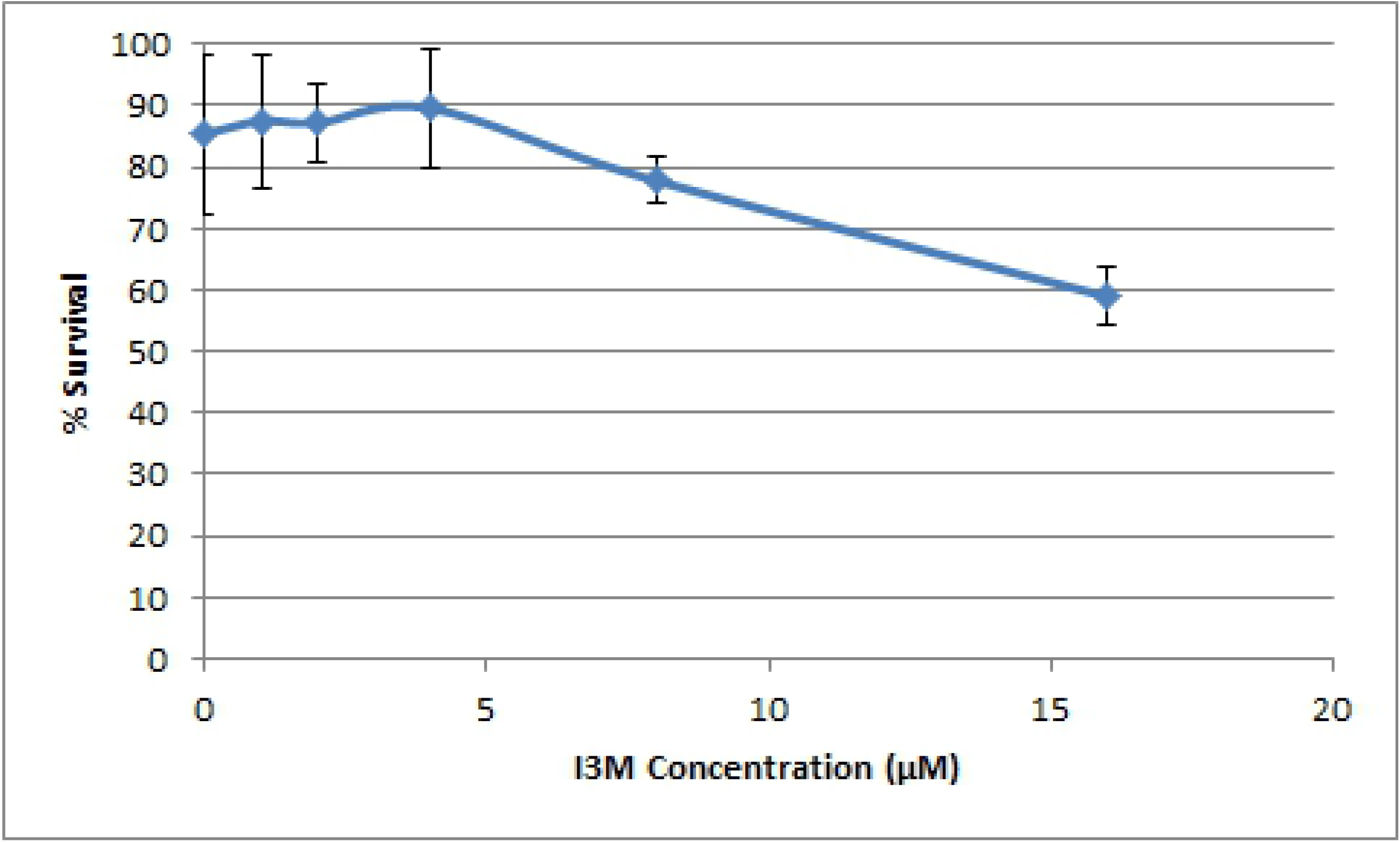
Survival curve for Tg(flk1:EGFP) zebrafish treated with various concentrations of I3M. The experiment was repeated in triplicate. The error bars represent the standard deviation of the survival percentage from the three experiments.

## 3 Discussion

This manuscript exhibits a functional high throughput drug discovery protocol to be used with zebrafish models. Using zebrafish in a high throughput drug discovery setting allows for an analysis of the effect of compounds on angiogenesis in a whole animal model *in vivo*. There are several transgenic zebrafish lines, such as the Tg(flk1:EGFP) line used in this work, and other models that allow for the visualization and quantification of angiogenesis [15]. In zebrafish, during angiogenesis TC and stalk cells (SC) are regulated by VEGF and Notch pathways, much like in human tumors [16]. This together with its high fecundity makes it ideal for statistically significant whole animal high throughput screening for angiogenesis inhibitors. The HTP provides a powerful and efficient tool for the discovery of drugs that modulate angiogenesis.

It is worthy to note that, in addition to the described methods, certain interventions were implemented to circumvent some limitations of working with the HTP and zebrafish. When dispensing with the COPAS there were a few rare occurrences where embryos would stick together or debris would be mistaken for an embryo. This would result in wells with errors (ie. no fish or more than one fish). If this was noticed during dispensing the protocol was paused and the wells with errors were corrected manually. In addition, due to the significant 4 day incubation time between the introduction of the compounds and imaging preparations, there was an opportunity to manually inspect the sample plates an hour or two just before imaging. Plates could be inspected for unhatched embryos that could be manually dechorionated and for visibly dead embryos. The visibly dead embryos usually coincided with a pixel count readout of less than 4000, which validated the use of that metric. Also, before imaging the sample plates were centrifuged, Centrifuge 5810 R [Eppendorf Canada, Mississauga, Ontario, Canada]. The centrifuge was ramped up to 1400 rpm at a rate of 140 rpm per second, then immediately back down at the same rate to force out any bubbles and encourage the fish to lie laterally on the bottom of the plate.

A potential use of the methods displayed herein could be to screen libraries of compounds that have already been through the rigors of clinical trials. This is highly significant because it is temporally and financially economical to use compounds already approved for clinical trials as repurposed drugs targeting pathological angiogenesis in the treatment of cancer. A main goal is to discover anti-angiogenesis compounds that target multiple or alternate mechanisms in tumor angiogenesis. This is important because cancers are known to promote angiogenesis via multiple pathways and not all cancers utilize these pathways in the same manner, with the different factors tipping the pro/anti-angiogenesis balance towards a pro-angiogenesis environment [17]. For example, there have been cases in which VEGF inhibitors were found to be more effective by also targeting other angiogenesis pathways in certain cancers [10]. The use of these methods is expected to result in the discovery of potentially novel anti-angiogenesis compounds that may improve treatment regimens for cancer.

The methods presented are from the perspective of a search for anti-angiogenesis compounds, but they can be adapted for other uses. The general order of steps in the methods would remain the same and similar pixel count readouts could be used for quantification in assays where a fluorescent readout is possible. The methods would simply need to consider certain key parameters for the best results: 1. Embryos would need to be dispensed after 8 hpf and ideally still be in their chorion for consistent time of flight (TOF) readings. Once out of the chorion, anesthetic will be required to keep the fish immobile for readings and fish orientation may vary, making it difficult to define a TOF window. 2. The time of compound introduction and length of incubation would be specific to the physiological phenomena being observed. 3. Acquiring readouts would be ideal at 3 or 4 dpf. The fish are more likely to have hatched and consistently lie laterally at these times. After 4 dpf, achieving a consistent orientation for imaging is less likely, due to the developing swim bladder.

## 4 Materials and methods

### 4.1 STX220 Incubator

A drug screen for endothelial cell modulators with the HTP begins with an empty 96-well optical bottom black wall plate in the incubator. This is done so that a rack position can be logged and associated with a specific plate. This allows data to be linked to a specific plate when there is an experiment that involves many plates. Alternatively, a bar code reader next to the incubator can be used for plates equipped with barcode. The incubator is also used to house sample plates after the compounds have been introduced. It was found that the condition of one embryo per well in a 96-well optical bottom black wall plate with 100 μL in each well created an environment where the embryos matured slower than expected. For this reason, the incubator was kept at 32 °C to encourage development and hatching by 4 dpf.

### 4.2 COPAS XL Embryo Sorter

The COPAS embryo sorter was used to dispense 1 fish/well into the 96-well plates at around 8 hpf. The embryos were destroyed if dispensing was attempted before 8 hpf. The Tg(Flk1:EGFP) embryos were selected and dispensed based on their TOF readings. Embryos were selected from a homozygous population; therefore, there was no need to sort according to fluorescence. To allow for the dispensing of 1 embryo/well the concentration of embryos in the sample cup had to be limited to avoid having multiple embryos dispensed at once. The average specific gravity of the embryos did not allow them to rise above a certain level while they were mixed with the systems magnetic stir bar. For this reason, the optimal embryo to E2 water levels were 100 embryos for a water level at 1.8 cm above the sample cup base, 200 embryos for water levels between 1.8 cm and 3.2 cm, and 300 embryos for levels above 3.2 cm. Adding more than 300 embryos would result in dispensing errors. In addition, a vigorous trial and error process was used to find the optimal settings to dispense 1 embryo/well with 40 μL of E2 embryo medium. The pressure parameters can be found in Figure 6 (Fig 6). All other settings can be found in Supplementary Figure 2 (S2 Fig). Supplementary Video 1 (S1 Vid) shows the COPAS in the process of dispensing 1 embryo/well in a 96-well plate. The markings at 1.8 cm and 3.2 cm above the sample cup base can also be observed in the video.

**Fig 6.**
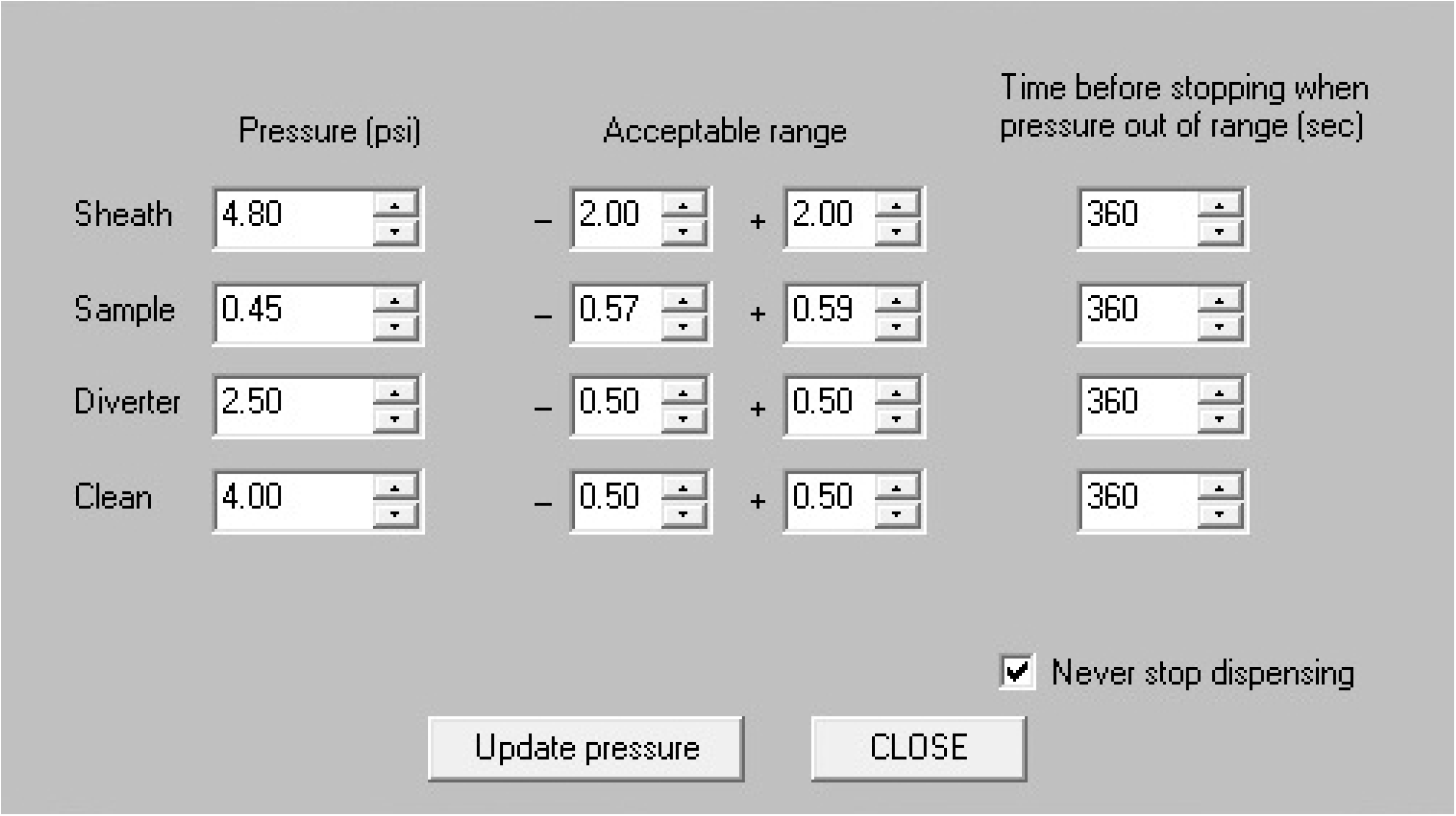
Optimized COPAS pressure settings to dispense 1 embryo/well. The “Never stop dispensing” box is checked so that dispensing does not stop due to the assigned pressures falling out of the default range.

### 4.3 Multidrop 384

The Multidrop was used to add 50 μL/well of a PTU/E2 media solution. A straight forward program was written to have the Multidrop dispense 50 μL/well. The Multidrop is triggered to dispense 50 μL/well once a 96-well plate is placed on it by the Twister II robot. Supplementary Figure 3 (S3 Fig) is a screen capture of the simple method that is triggered.

### 4.4 Sciclone G3 Advanced Liquid Handler

The Sciclone was used to dispense compounds from the compound plate into the sample plate. The application for this step was written in the Maestro Software provided with the Sciclone from Caliper Life Sciences. This program begins with initializing the Sciclone, homing the X, Y, and Z axes of the head device, and setting the movement speed to 100%. The system then loads 200 μL non-filtered and non-sterile automation pipet tips [Caliper] onto the head device from a tip box. The head device then moves to the compound plate and aspirates 10 μL as a trailing air gap to ensure that the entirety of the compound is dispensed later on. The head device is set to descend to a position where each tip is positioned 2 mm above the bottom of its corresponding well. Here 10 μL of compound per well is aspirated at 30 μL/s. The head then retracts at 10% speed, to not leave any residual compound on the outside of the tips, and aspirates 10 μL as a leading air gap, to prevent any compound from leaking out of the tips. The head then moves to the sample plate and descends to have the tips 2 mm above the bottom of their corresponding well and all of the tip content is dispensed. The head device then moves above a waste receptacle and dispenses the used tips. From here grips on the head device pick up the sample plate and move it to a shaker. Here the plate is shaken in a figure eight pattern at 100 rpm for 10 seconds. The plate is then moved back to its original position. The head device is then brought back to the home position to make way for the twister II robot that brings the sample plate to the incubator. Supplementary Video 2 (S2 Vid) shows the Sciclone acting out this protocol. Supplementary Figure 4 (S4 Fig) shows the application script.

The Sciclone was also used to dispense clove oil from a reservoir container into the sample plate. The program used for this is identical the one described for dispensing compounds into the sample plate from the compound plate, with one difference being that the tips descend into a reservoir container at 0.2 mm above its bottom for aspiration instead of 2 mm above the bottom of a 96-well plate.

### 4.5 Image Xpress Ultra

The Image Xpress Ultra confocal imager was used to capture a 2D image of the Tg(Flk1:EGFP) zebrafish in each well of the 96-well plate at 4 dpf. A 4x objective was used to capture the whole well and thus, ensure that the entirety of each fish was imaged. Each well was imaged in four evenly sized quadrants at four times averaging, with a scan size of 2000 x 2000 pixels, to accomplish this. The z-offset from the bottom of the wells was optimal at −0.3 μm. These settings resulted in 384 images per plate, requiring 733.9 MB of storage. Each set of 4 images per well took roughly 15 s to acquire. A journal was developed in Meta Xpress to process the images from each well. The journal begins by stitching the 2×2 images from each well into one image with a 10% overlap between the images to ensure proper alignment. Each image is then processed with a pixel counting program. This program identifies and counts pixels representing endothelial tissue using a lower pixel intensity threshold of 3000 on the software arbitrary scale. This allows for pixels representing endothelial tissue in the focal plane and out of the focal plane to be included, while excluding background data. Thus, allowing the data from a 2D image to be more representative of a 3D object. This type of processing removes the need to work with stacks of images to create 3D representations and in turn significantly reduces imaging and processing time to a level more favourable for a high throughput process. These Image Express Ultra settings and the Meta Xpress journal were also used in previous work describing a zebrafish sepsis model for high throughput drug discovery but not described in detail [18].

### 4.6 Zebrafish

The Tg(Flk1:EGFP) line was maintained and crossed using standard techniques [19]. All zebrafish experiments were approved by the St. Michael’s Hospital Animal Care Committee (Toronto, Ontario, Canada) under protocol ACC867.

### 4.7 Code Availability

All efforts were made to include all parameters and data required for anyone to repeat these methods or adapt them for use in other types of assays. In addition to this, sample code and further details can be found amongst the supplementary images. Any journal that was developed in Meta Xpress was a collaborative effort with Molecular Devices and can be obtained from the corresponding author upon request or directly from a Molecular Devices application scientist.

## 5 Data availability statement

The data generated and/or analysed during this work, if not already included in this manuscript, can be obtained from the corresponding author upon request.

## Supporting information

S1_Vid

S2_Vid

## Acknowledgements

We thank the technical staff at Caliper (a division of PerkinElmer) [Hopkinton, Massachusetts, USA, Liconic Instruments [Somerset, New Jersey, USA], Union Biometrica [Holliston, Massachusetts, USA], Thermo Scientific [Hudson, New Hampshire, USA], and Molecular Devices [Sunnyvale, California, USA] for aiding in the construction and functional development of the HTP. This work was supported by Canada Foundation for Innovation (grant #26233) (X.Y.W.) and Ontario Centres of Excellence-Consortium Québécois sur la Découverte du Médicament (OCE-CQDM) Life Sciences R&D Challenge Program (X.Y.W).

## Author Contributions

Conceptualization: AM, XYW.

Data Curation: AM.

Formal Analysis: AM.

Funding Acquisition: XYW.

Investigation: AM.

Methodology: AM.

Project Administration: AM, XYW.

Resources: YW, XYW.

Software: AM.

Supervision: AM, KKS, XYW.

Validation: AM, RN Visualisation: AM, RN, JYL.

Writing – Original Draft Preparation: AM.

Writing – Review & Editing: AM, RN, RG, KKS, XYW

## Supporting Information Captions Supplementary Figures

**S1 Fig** – **Master iLink Method used to instruct the HTS to execute the protocol to discover angiogenesis inhibiting compounds.** The method begins by initializing all the required equipment and confirming that all plates and consumable are programed in a logical location. As the robots place the 96-well sample plates on the device corresponding to a certain protocol step a separate routine, program, protocol, or simple instruction is triggered by iLink and run by the device. When the device is done working through the set of commands, it triggers iLink to move onto the next step.

**S2 Fig – Optimized COPAS settings to dispense 1 embryo/well.**

**S3 Fig – Screen shot of the simple Multidrop method triggered when a 96-well sample plate is placed on it.**

**S4 Fig – Application script used to have the Sciclone liquid handler dispense 10 μL of compound from each well of a 96-well compound plate into each well of a 96-well sample plate.**

## Supplementary Videos

**S1 Vid - COPAS in the process of dispensing 1 embryo/well in a 96-well plate.**

**S2 Vid - Sciclone acting out protocol to aspirate compounds from the compound plate and dispense them into the sample plate.**

