## Supplementary figures and images for "Protocol development for discovery of angiogenesis inhibitors *via* automated methods using zebrafish"

### S1_Fig

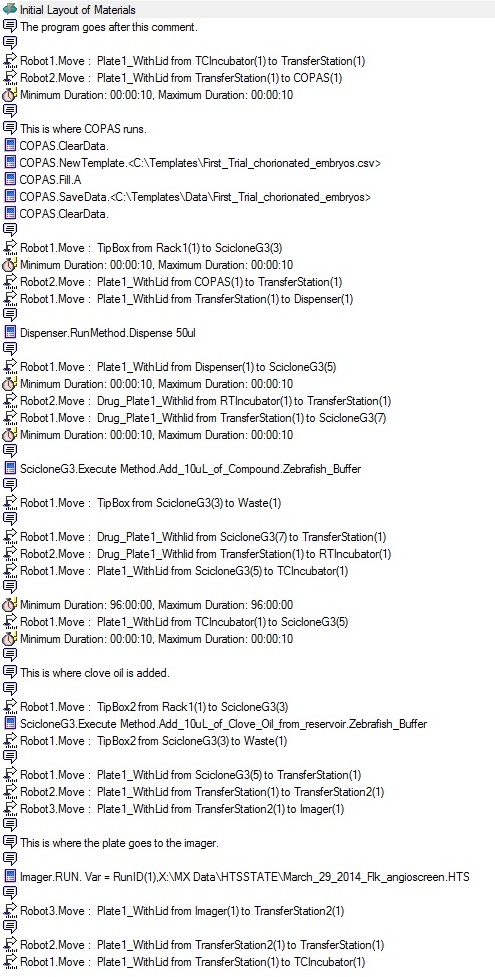

### S2_Fig

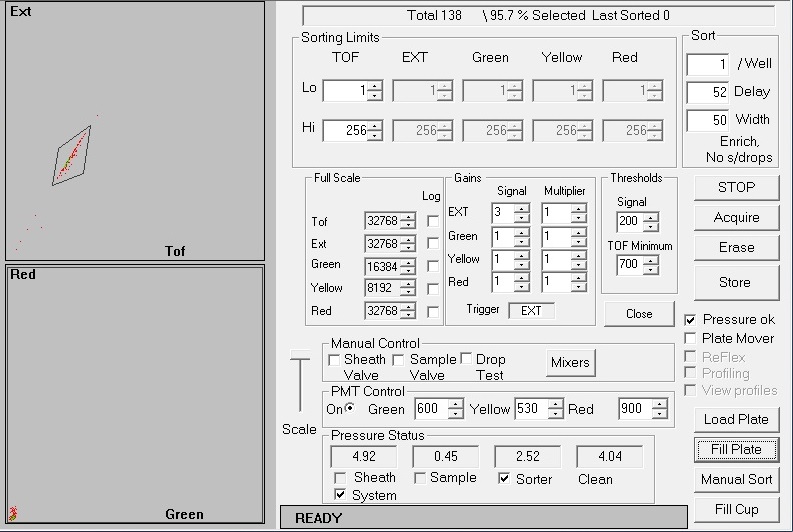

### S3_Fig

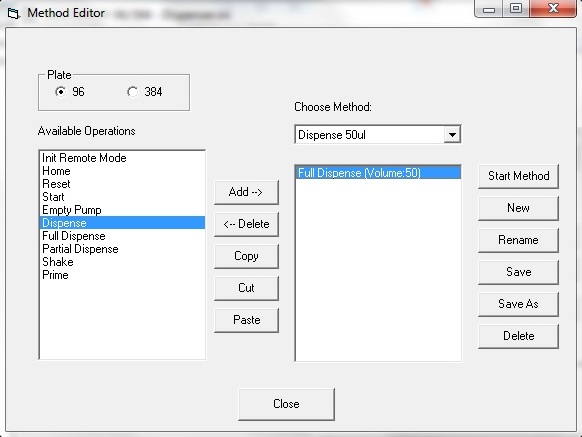

### S4_Fig

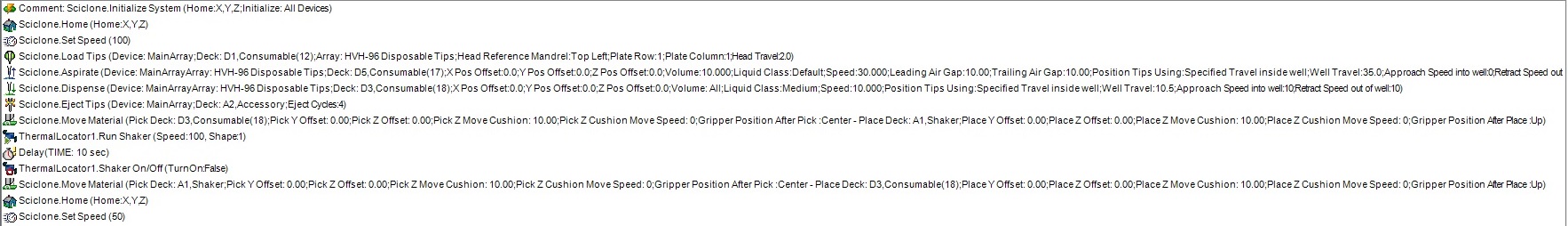
